# Survey of Core Facilities shows the importance of communication and management for optimal research quality

**DOI:** 10.1101/2020.08.19.256511

**Authors:** IC Kos-Braun, B Gerlach, C Pitzer

## Abstract

Recently, it has become evident that academic research faces issues with the reproducibility of research data. It is critical to understand the underlying causes in order to remedy this situation. Core Facilities (CFs) have a central position in the research infrastructure and therefore they are ideally suited to promote and disseminate good research standards through their users. However, there are currently no clear guidelines directly applicable to academic CFs. To identify the most important factors for research quality, we polled 253 CFs across Europe about their practices and analysed in detail the interaction process between CFs and their users, from the first contact to the publication of the results. Although the survey showed that CFs are dedicated to train and advise their users, it highlighted the following areas, the improvement of which would directly increase research quality: 1) motivating users to follow the advice and procedures for best research practice, 2) providing clear guidance on data management practices, 3) improving communication along the whole research process and 4) clearly defining the responsibilities of each party.

## Introduction

Academic life science research faces issues with the irreproducibility of research data. Many scientific studies are difficult or impossible to replicate (Ioannidis 2005; Prinz, Schlange, and Asadullah 2011; Ioannidis et al. 2014; Eisner 2018). A Nature survey asked 1,576 scientists for their thoughts on reproducibility (Baker 2016). Most researchers agree that there is a “significant crisis” and over 70% said they “have tried and failed to reproduce another scientist’s experiments”. There are several causes for this situation: selective reporting to support favourite hypothesis, pressure to publish, poor analysis, lack of replication and unexperienced or insufficiently trained researchers (Baker 2016; Smaldino and McElreath, 2016).

The Association of Biomolecular Resource Facilities (ABRF) surveyed over 200 Core Facilities (CFs) to assess how they implemented the U.S. National Institutes of Health (NIH) guidelines on scientific rigor and reproducibility (Knudtson et al. 2019). The survey identified sample quality as the major challenge. In addition, the lack of training, expertise or supervision were the main factors affecting the quality of research.

These observations show that the scientific community is increasingly acknowledging data integrity as a major issue and trying to remedy it by setting up several initiatives and promoting guidelines for good research practices (Munafò et al. 2017). One of these initiatives is the Centre for Open Science (COS) which also published the “TOP-guidelines” (TOP = Transparency and Openness Promotion) in 2015 aiming for an increased transparency of the research process and the preregistration of experiments (Nosek et al. 2015). An important discussion is currently arising about quality systems and auditing for academic research (Dirnagl et al. 2018). This is facilitated by the development of a quality system for biomedical research by the European Quality in Preclinical Data consortium (EQIPD). It provides a user-friendly, lean and flexible approach to increase quality of research data that can be implemented in any research unit (Bespalov et al. 2020). FAIR (Wilkinson et al. 2016) and ALCOAplus can be named as two among several best practices specifically aiming to achieve better data integrity by setting standards for documentation of meta data and describing information for data integrity. Another initiative is Reproducibility2020 from the Global Biological Standards Institute (GBSI) which addresses the concern of irreproducible biomedical research (Freedman, Venugopalan, and Wisman 2017).

CFs have a central position in academic life science research: they provide access to state-of-the-art equipment and advanced skills in a cost and time effective way. They develop new technologies and spread their technical and research expertise to the numerous scientists they teach. They connect institutions, foster exchange, collaborations and interdisciplinary research (Meder et al. 2016). CFs generate a substantial part of the scientific data at academic life science research institutions. They offer a protection against bias in experimental design and data analysis, therefore support transparency, rigor and reproducibility. In addition, due to their central position in the research infrastructure, CFs are ideally suited to promote quality standards and disseminate good laboratory practice (GLP) through training and workshops. The impact of CFs on research culture is broad and long lasting, as they also teach many young researchers who can then continue to spread the acquired habits during their careers.

Therefore, we assessed in detail how CFs implement good research practices and transmit them to their users. We asked CFs from different fields of life sciences across Europe about their practices, in order to analyse the whole process of data generation from the first contact with the researcher to the publication of results. Our survey shows that CFs are dedicated in providing the best possible service to their customers, despite the fact that half of them struggle with insufficient funding. We uncovered issues that affect the quality of research at CFs, such as the lack of quality checkpoints and the importance of a good facility management system, an efficient communication with users and a clear definition of responsibilities.

## Results

Our survey was sent to 1000 CFs from different fields of life sciences throughout Europe. We received 253 complete forms from over 30 types of CFs, which differ in the techniques and expertise they offer (Figure S1a). They vary in the number of employees and users, as well as the amount of generated data (Figure S1b - d).

### Core facilities are dedicated to service

We assessed CFs’ quality procedures and interactions with the users during the whole research process from experimental design to publication. The CFs were asked several yes/no questions about quality practices (Figure 1). Overall, the results show that the CFs are dedicated to providing the best possible service to their users. The majority of CFs offer training and guidance on experimental design, sample preparation, data analysis and help troubleshooting. The also offer support in writing relevant sections of publications. At the same time, the survey identified weaker areas with potential for improvement, such as communication with users and management, which we will examine in more detail below.

**Fig 1.**
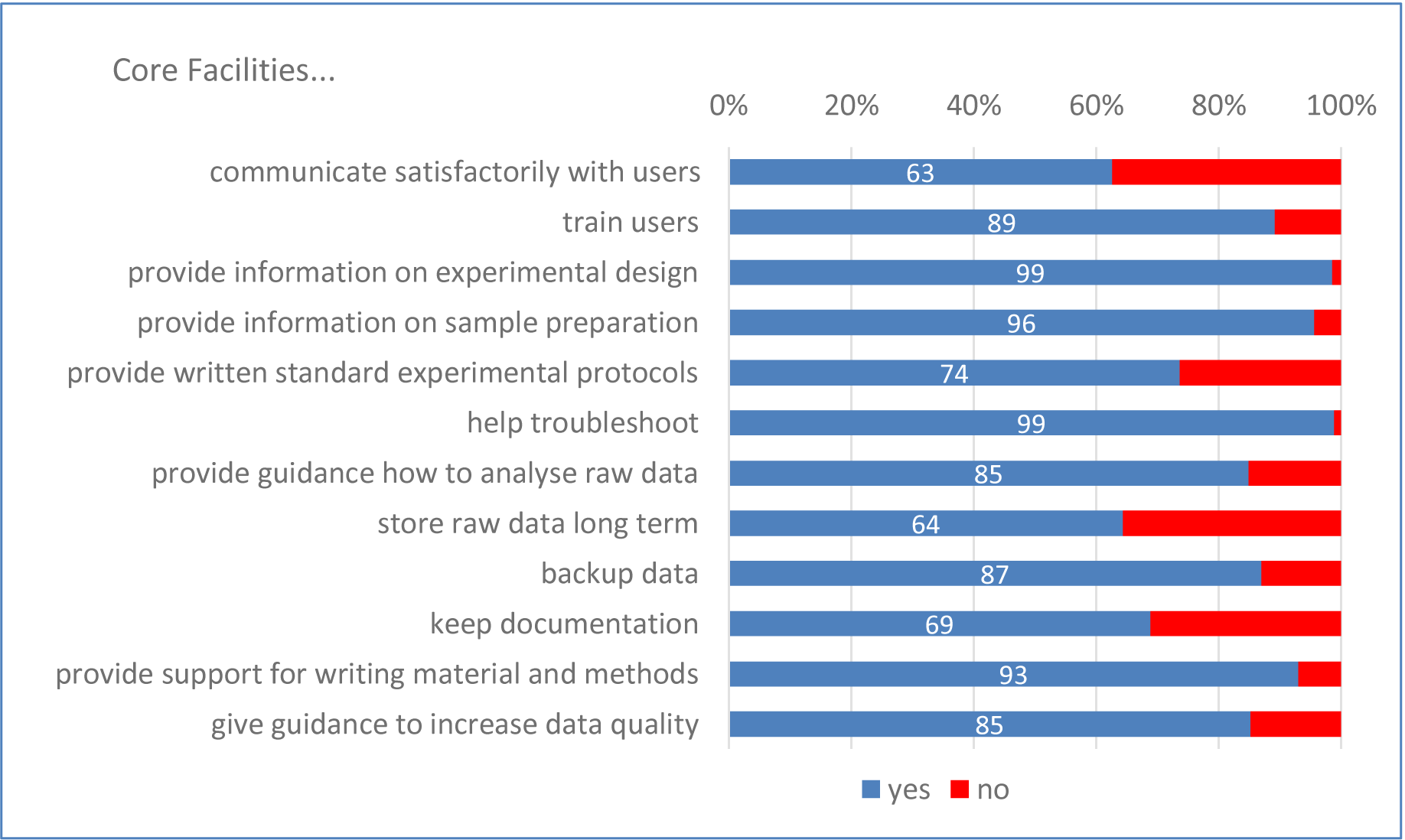
Services provided by the core facilities along the research process in percentage.

### Full-service versus self-service facilities

CFs can be classified in three different groups depending on who performs the experiment at the facility: 1) the CF staff, 2) the external researcher (user) or 3) both CF staff and users. We call *full-service* facilities the CFs offering an “all-inclusive service”, where the CF staff executes the experiment with or without data analysis. Other CFs, called “user laboratories” in (Meder et al. 2016), provide and maintain an infrastructure where users have access to equipment, training and expert advice. Here we refer to these as *self-service* facilities. The last group, which we call the *hybrid-service* facilities, is a mixture from both previous operating modes where both staff and users are performing experiments.

In addition to the distinction of who actually performs the experiments, the service range provided by the CFs varies as well, from a basic one consisting of processing the samples and sending back the data to an extended range from experimental design to publication. However, we do not specifically distinguish between these options.

Most CFs in our survey belong to the hybrid-service facilities, followed by full-service ones, while self-service facilities are the least common (Figure 2a). Although most aspects are similar in all three types of CFs, there are notable differences between the operating modes (Figure 2b). Only about a quarter of self-service facilities keeps documentation of the experiments compared to >95% of full-service CFs. Similarly, storage of raw data is offered by only half as many self-service CFs (40%) compared to full-service CFs (82%). Furthermore, fewer self-service CFs provide standard experimental protocols because the users may bring their own.

**Fig 2.**
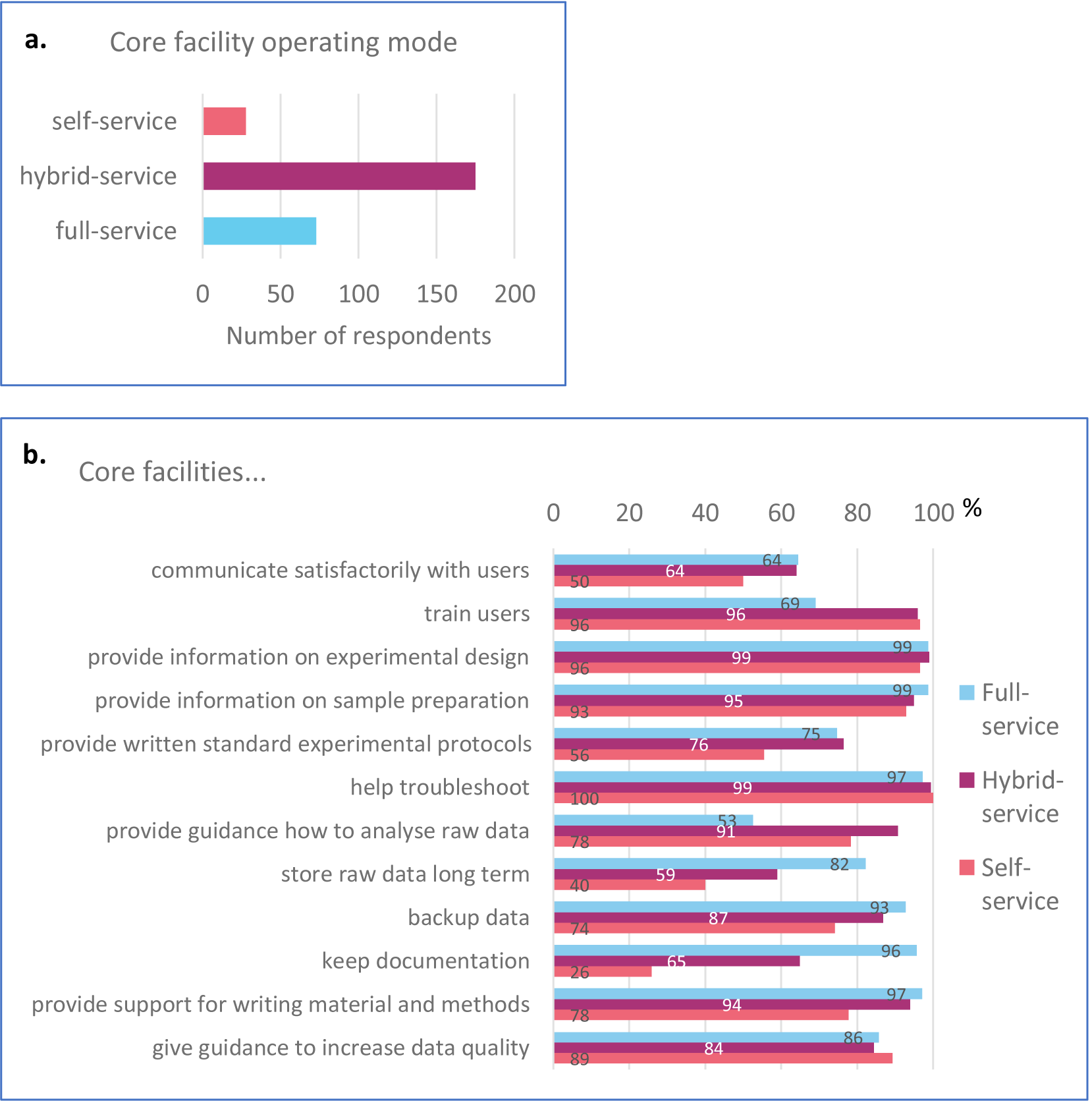
Full-service versus self-service facilities. a) Number of Core facilities by operating modes. b) Services provided by the core facilities along the research process depending on their mode

On the other hand, the full-service CFs tend to train their users less, consider training less important and provide primarily theoretical training (Figure 2b and S2a, b). Only half of full-service CFs provide guidance how to analyse raw data, because they analyse the data themselves (Figure 2b and S2c).

### Research quality

#### Lack of funding is the major obstacle to research quality

In order to identify what is critical for research quality, we asked CFs an open-field text question to list what factors they consider the most important and which of these need to be improved at their facility. As can be seen from the Figure 3a, the factors considered the most important are training and communicating with users, followed by having enough qualified staff, as well as up-to-date and well-maintained equipment. From these factors, hiring more staff and purchasing/maintaining equipment are the most in need of improvement. These are also the costliest ones. Interestingly, although not considered as important, the aspect listed second in need of improvement (directly after hiring more staff) is management. Management was mentioned at many different levels: facility, projects, samples, data, IT infrastructure, documentation or automation (discussed in more detail below).

**Fig 3.**
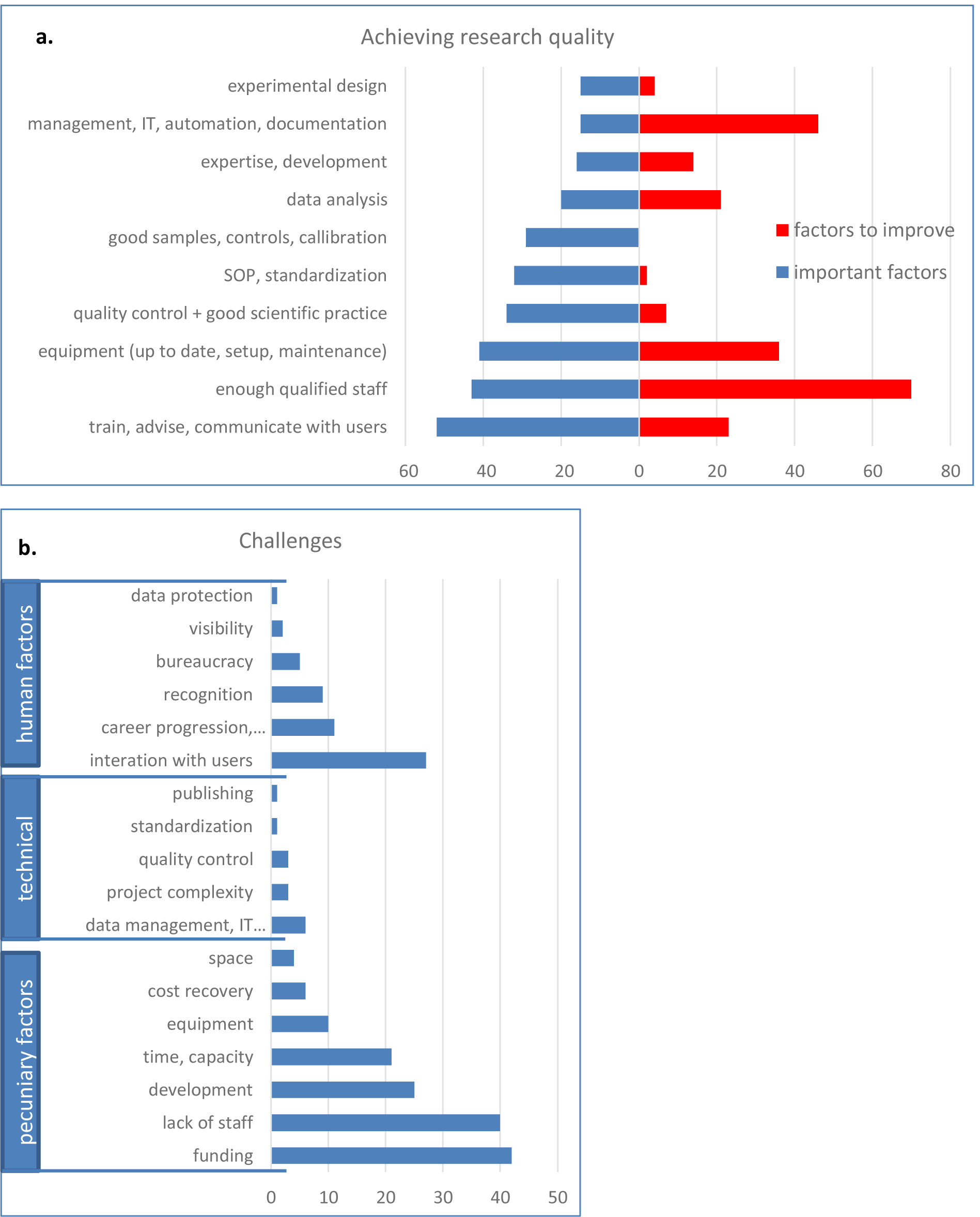
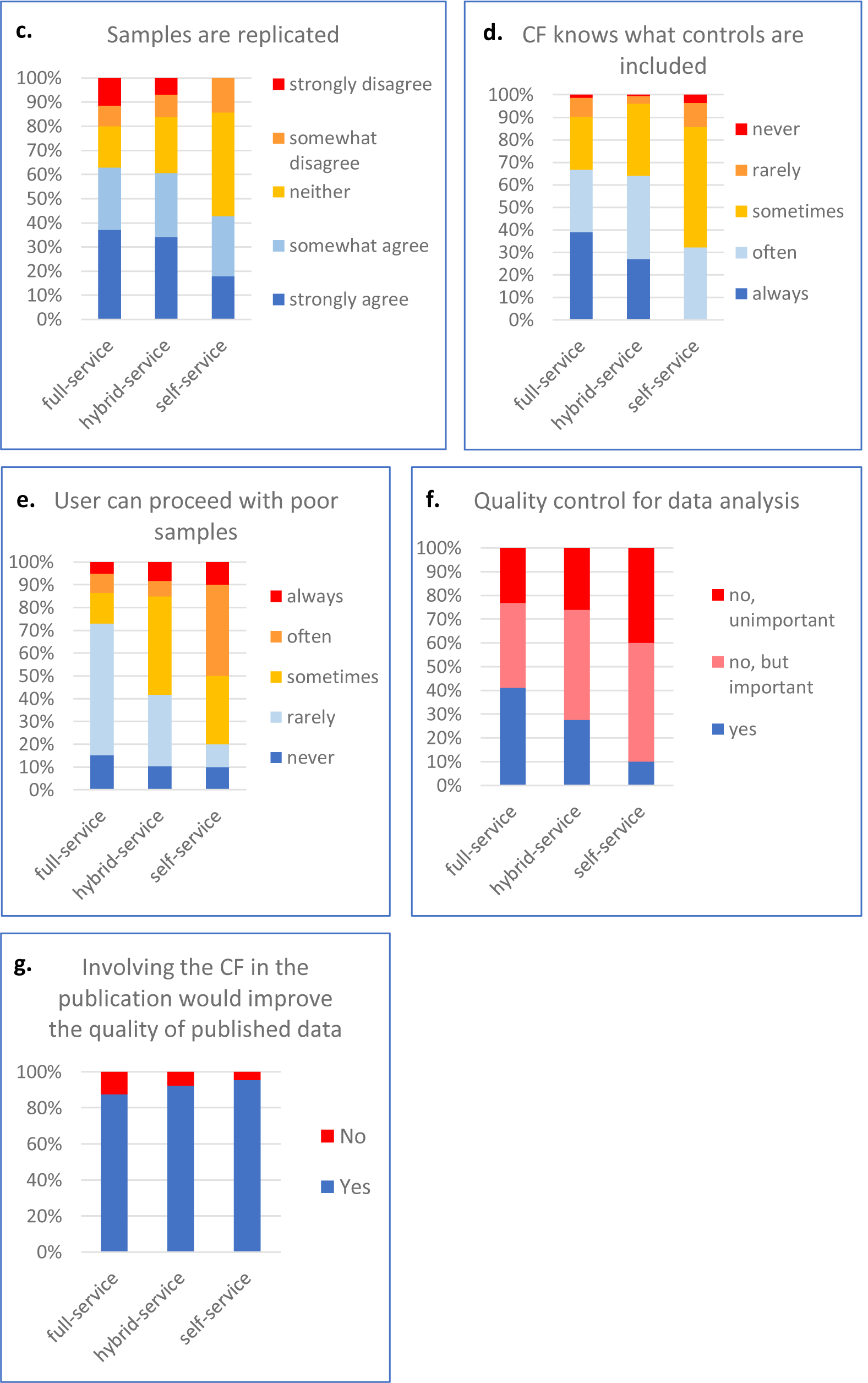
Research quality. a) The most important factors for achieving research quality (in blue) and the ones that need to be improved (in red). b) Challenges faced by CFs. The category “career progression” includes “permanent positions” and “motivation”. IT means “IT infrastructure”. c) Samples replication. d) Controls inclusion. e) Percentage of CFs allowing users to proceed with poor samples. f) Importance of quality control for data analysis g) Importance of involving the CFs in the publication to improve the quality of published data

When asked about the biggest challenges encountered by CFs, the most frequently cited was unsurprisingly the lack of funding, closely followed by the lack of staff (Figure 3b). Frequently cited were also development (keeping the facility efficient and state-of-the-art), time and capacity. All these challenges arise in fact from the lack of funding. About half CFs do not have enough staff and/or enough funding (data not shown). The third most frequently quoted challenge is not related to funding but concerns the interaction of CFs’ staff with the users (discussed in more detail below).

#### Controlling quality from experimental design to publication

The process from experimental design to publication can be controlled at multiple checkpoints. As already shown in the Figure 1, CFs strive to provide guidance along the whole process, from the experimental design, sample preparation, experimental protocol (SOPs) to data analysis.

Starting with the experimental design – how many samples, which controls should be included, how the experiment will be performed and evaluated – all these points should be discussed with and use the expertise of the CF staff. Surprisingly, experiments are often performed without replication, which is essential for good quality research (Figure 3c). Even when it comes experimental controls, CFs often do not know what controls were included (Figure 3d), even though experiments lacking appropriate controls cannot be meaningfully evaluated.

When asked if users could proceed with poor samples, we found that 50% of self-service facilities often or always allow users to analyse poor samples. Furthermore, even in full-service CFs, 13% of these always or often accepts poor samples (Figure 3e). Regardless of the reason, whether it is due to the lack of controlling the sample quality or knowingly accepting poor samples, the use of such samples counteracts the efforts of achieve good quality research. Of course, such samples can be occasionally justified in special cases, but it should not be the norm. Introducing a sample quality checkpoint before the start of experiment is a simple measure that would clearly increase the quality of produced data.

Regarding data analysis, 40% of full-service CFs and only 10% of self-service CFs have mechanisms to ensure correct analysis and interpretation of raw data (Figure 3f). Data analysis quality control is usually performed by having the data checked by another CF staff member. It can also involve discussing the results with the user or the PI, or using internal standards and quality control samples (data not shown). While the majority of CFs do not control the quality of data analysis, most of them consider it important to have (Figure 3f). It is however often difficult to implement mostly due to the lack of staff.

The last opportunity for CFs to check if the data they helped to produce was analysed, interpreted and presented correctly is before publication. However, CFs are often not even informed about the upcoming publication (see the section on communication below). The vast majority (91%) of all CFs believe that if they were involved in the publication process it would improve the quality of published data (Figure 3g). This is most important in self-service CFs. The following quotes from different participants illustrate the opinion on or experience with the CFs involvement in publication process:

- “[It] ensures correct understanding and an accurate account of what happened”
- “The users often lack the knowledge to use the correct controls or ways of display, without being aware that they are not following best-practice.”
- “The core facility can ensure that the methods are detailed so that they can be replicated.”

In conclusion, it is pertinent to introduce checkpoints to control the experimental design, sample quality, data analysis, and methods section and figures for publication. This seems particularly important in self-service facilities where more supervision would benefit the research quality.

### Management

Management is a very important factor for achieving research quality and many CFs recognised the need to improve it (Figure 3a). Obviously, there are many levels of management, such as managing the budget, users, projects, samples and generated data.

A management software is the most frequently used tool by CFs to achieve research quality (Figure S3a). Overall, close to 30% CFs use a management software and further 34% believe it would make sense to use one (Figure 4a).

**Fig 4.**
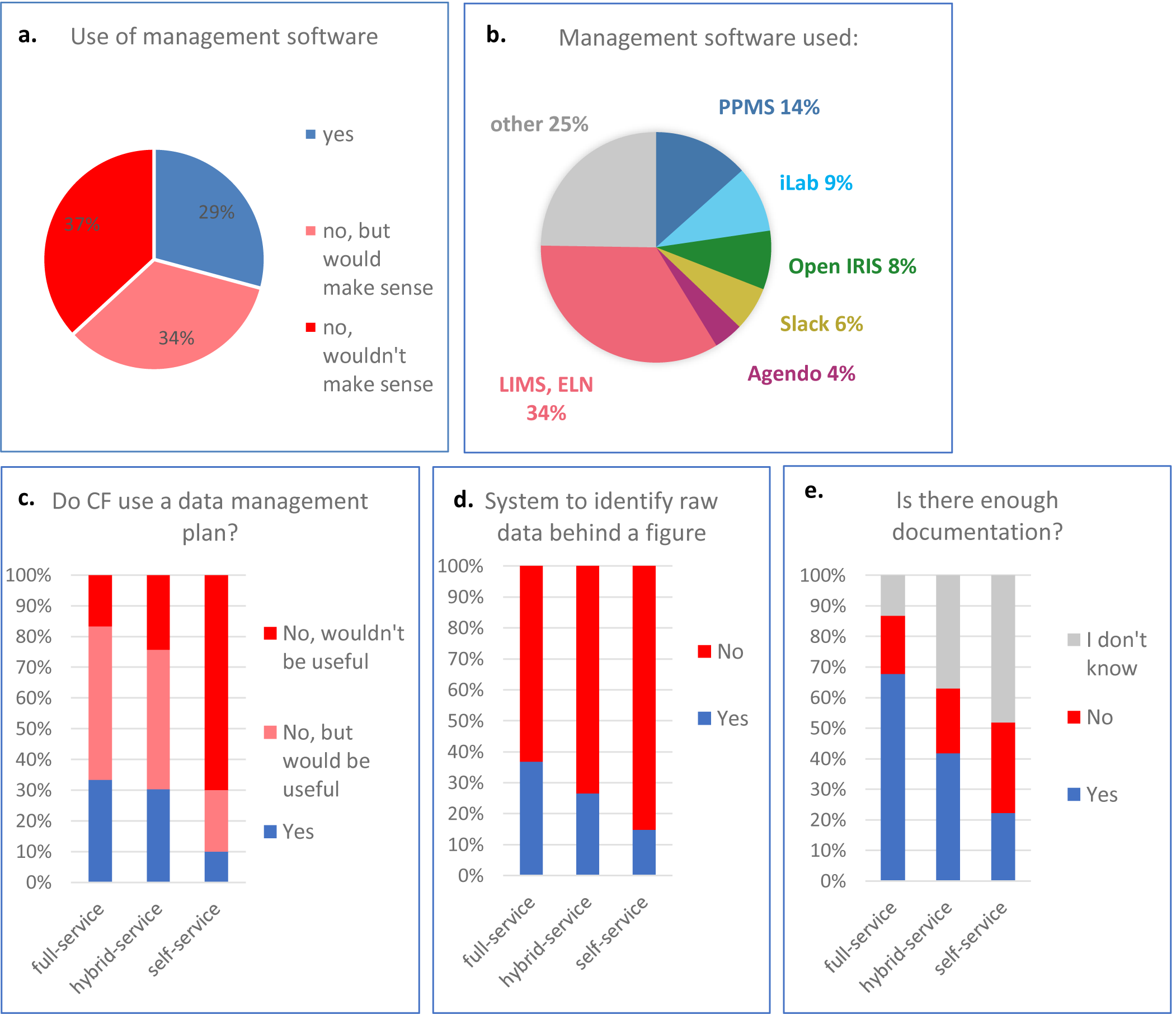
Management. a) Proportion of CFs using management software b) Management software used. “Other” is software used by only 1 or 2 CF in the survey c) Proportion of CFs using a data management plan d) Proportion of CFs having a system to identify raw data behind a published figure e) Documentation in different CF types

Currently, the management software used in CFs can be split into two categories (Figure S3b). The first category, “core facility managements software” mainly allow CFs to communicate with their users, book equipment and manage access rights, training, maintenance, technical issues and billing. It can also manage individual projects to a certain extent and keep records in the form of uploaded documents. Examples of such CF management software are PPMS from Stratocore, iLab from Agilent, Agendo or Open IRIS (open source). The second category is the “data management software”. Electronic Lab Notebooks (ELN) allow the precise recording of scientific procedures from the experimental design and sample preparation to the publication. It manages data acquisition, storage and analysis. This interconnected documentation ensures transparency and traceability. Laboratory Information Management Systems (LIMS) are similar although they are often linked to one piece of equipment.

Our survey revealed that the software solutions used by CFs are very heterogeneous (Figure 4b). About 35% CFs are using CF management software (PPMS, iLab, Open IRIS or Agendo) and 35% CFs are using data management software (LIMS or ELN). 4% CFs are using both. From the CF management software, PPMS is the mostly used one in self-service facilities, while iLab, open IRIS and Agendo dominate in full-service CFs (Figure S3c). Notably, a quarter of CFs uses software solutions mentioned only once or twice in our survey (“other” in Figure S3c). The data management software is used in all types of CFs to a similar extent.

However, CFs also mentioned drawbacks in using a specialized software, such as lack of cooperation of users and difficulty to customise for heterogeneous and often complex projects. The implementation and cost of such software were also considered a problem (data not shown).

We also asked if CFs have implemented a “data management plan” instructing how research data will be annotated, stored and analysed. A data management plan ensures that all data remain traceable. It is used in 30% full-service but only 10% self-service CFs (Figure 4c). Another 50% full-service CFs believe it would be useful, while only 20% self-service do.

Looking into the different aspects of data management, we saw that about half of CFs have implemented data management measures to ensure that data are complete, attributable, reusable, compatible, searchable and findable (Figure S3d).

Strikingly, 72% of all CFs do not have a system to identify raw data used for published figures (85% and 63% for self-service and full-service respectively) (Figure 4d). The remaining CFs mentioned that they use a data management software (ELN, LIMS), unique IDs or a public repository to trace raw data (data not shown). The problem of non-traceable data is linked to the issue of insufficient documentation of experiments, which is clearly more pronounced in self-service facilities (Figure 4e). Only 20% of self-service CFs have enough documentation, while 70% full-service CFs document their experiments sufficiently. Importantly, a half of self-service CFs do not actually know, how the experiments are documented. This might be connected to the issue of communication and responsibility, addressed below.

### Communication, respect and trust between CFs and users

Communication plays a critical role in the interaction between CFs and their users, as CFs provide a service based on their users’ requirements and users need to prepare their samples and experiments according to the advice of CFs staff who has expertise in the equipment and techniques available in their CF. Communication is regarded by CFs as a sensitive issue and the interaction with the users is seen as a challenge (Figure 3b). About 37% CFs feel that the communication with their users needs to be improved (Figure 5a). Communication between CFs staff and users is mostly done through emails and/or in person and CFs say it could be improved by using a communication or management software or a chat/discussion platform, and actively motivating the user to cooperate and read the information provided (data not shown).

**Fig 5.**
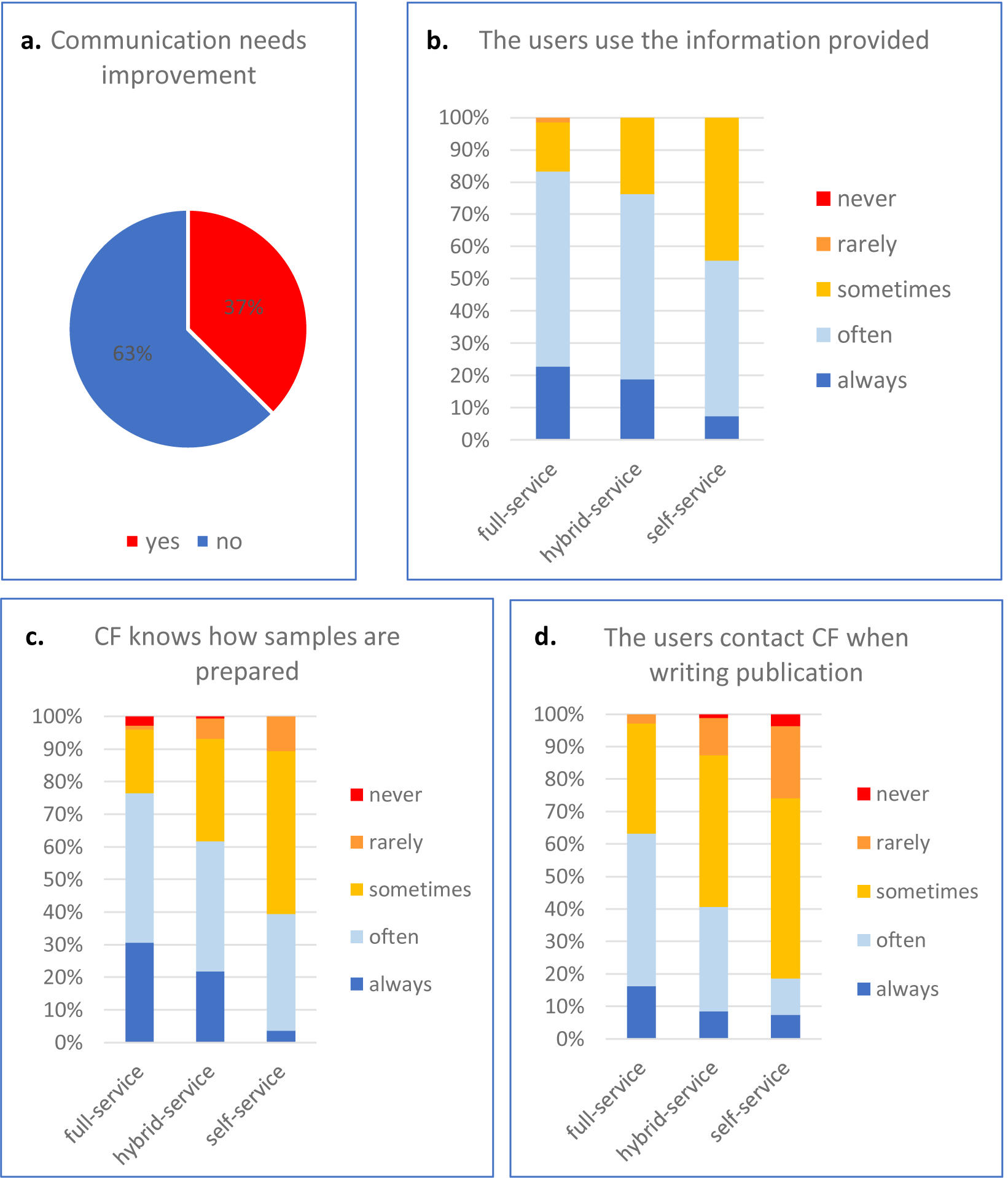
Communication and interaction between CFs and users. a) Does communication between the CF staff and their users need improvement? b) How the users use the information provided by the CF staff c) Awareness about sample preparation d) Contact between users and CF staff when writing a publication

Communication between staff and users is a common cause of tensions. The following selected comments from different participants illustrate the communication issue:

- “If I produce a plan, it will just be another formal document that will be ignored…”
- “It is very hard to get users to engage in this [quality control of data analysis]. It’s hard enough to get them to use the correct controls!”
- “One of the problems that we have had is scientists thinking they know how to do analysis and using the incorrect statistical test or website because it gives them the answer they were after rather than the correct answer.”
- “It is sometimes difficult - usually more because of group leaders than because of students - to get people to accept new or improved ways of doing certain types of experiments.”
- “The core facility tried to implement a data management plan, but this was not accepted by the user.”

Crucially, these quotes also reveal another important issue: many users (or their PIs) seem not to trust or respect the expertise of CFs personnel. Although CFs are committed to help their users and most of them (85 to 99%) provide information from experimental design to publication (Figure 1), there is a gap in the uptake from the user’s side. About half of self-service CFs estimate that their users use this information only rarely or just sometimes (Figure 5b). Users use the provided information more frequently in hybrid-service and full-service facilities, which is likely due to the need to conform to the facility’s specific instructions for sample preparation.

Furthermore, the communication/trust issue affects another aspect critical for good quality research, the evaluation of sample quality (also discussed in the section about the quality of research). Over 70% and 95% CFs (full-service and self-service respectively) do not have a full knowledge how samples are prepared (Figure 5c).

Finally, less than 20% users of self-service facilities contact the CFs (always or often) before publishing their results. CFs deplore it because they believe they could improve the quality of the published data (Figure 3e).

Together these results show that even though communication, trust and respect are sensitive topics, they nevertheless must be fostered as it is essential for good quality science.

### Sharing responsibility between CFs and their users

Unexpectedly, numerous answers to free text questions raised the issue of responsibility, although our survey did not specifically examine this aspect. The words “responsible” or “responsibility” were mentioned in total 123 times, referring to issues ranging from experimental design to publication. It is undeniable that responsibility is an essential aspect of good quality research. However, the quotes from CFs revealed an ambiguity in discerning “who is responsible for what”. Most CFs do not see themselves responsible, as one participant explained: “We allow poor samples to be processed, since the responsibility for the experiment lies entirely on the researcher!”. Another respondent stated: “The users are responsible for their data.” And one participant stressed: “We strongly feel that responsibility for data analysis and interpretation must be in hands of researchers, especially in the hands of research group leaders who are responsible for final research outcome.”

On the other hand, a small number of CFs do consider themselves responsible for the produced data quality, as one participant wrote that “the Core Facilities should be more involved in planning of the experiments and should be also responsible for the data generated in the Core Facility.” Another one explained that “it is the overall responsibility of the facility to make sure that data are analysed correctly. If a user decides to analyse their data, we will make sure at the publication stage that all data and conclusions drawn are consistent.” The responsibility for data quality can also be integrated into the internal rules: “It is the policy of our institute that all data generated through platforms is checked by the platform staff/head before publication”.

The lack of clarity in the responsibility rules can negatively affects the quality of research. As an example, CFs do not agree who should store the raw data, which is probably also one of the reasons for the lack of traceability (as presented in the section on data management). Many respondents believe that the “long-term [data] storage is a responsibility of each individual group leader”. The 30% CFs, which do not offer data storage, mostly believe they should not (Figure 6). Yet, one participant acknowledges the merit in storing raw data: “I think the responsibility for storing data and having back-ups is with the user. However, to have a backup of raw data at the core facility would possibly discourage users to perform improper data manipulation and could help to solve issues on scientific misconduct.”

**Fig 6.**
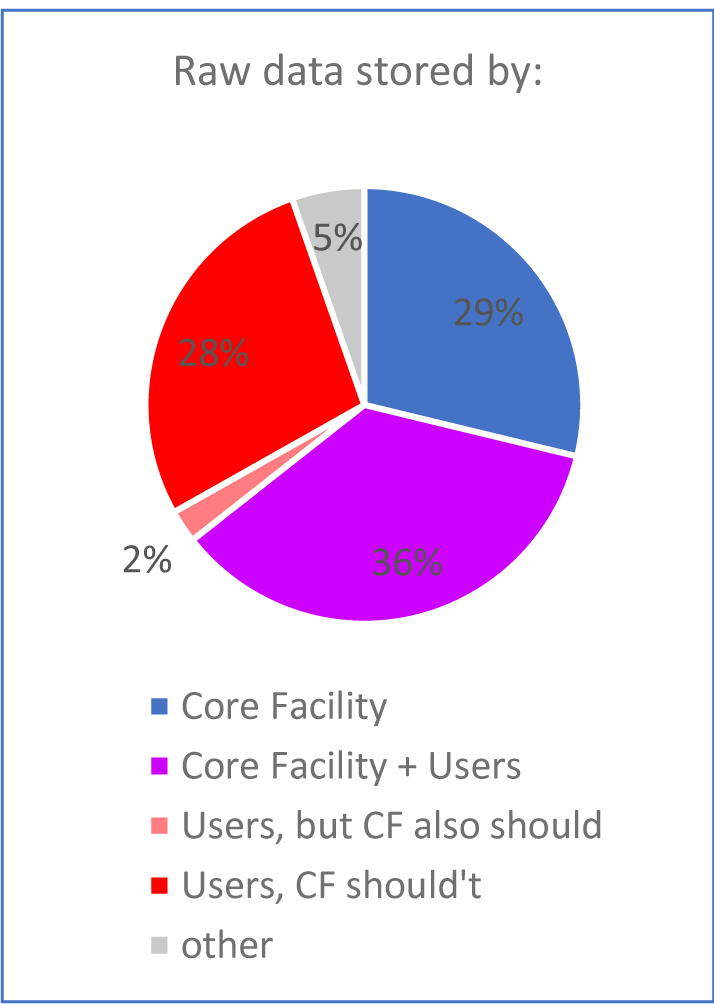
Responsibility for storing raw data.

Most of the time CFs are not listed as authors on publications. Contradictorily, the majority of CFs say they are not responsible for the generated data, but at the same time believe that being part of the publication process would improve the quality of the published data (Figure 3e). It is important to realize that responsibility cannot be simply off-loaded. All the parties involved in the experiments share responsibility for the data generated. This especially applies to all the authors on a publication, who all share responsibility for accuracy of the published data, as one participant wrote: “Being part of publication is holding responsibility for the work done. Neither the researchers, inclusively the PI’s are in a place to take responsibility for work with technologies that they do not understand.”

## Discussion

We surveyed 253 CFs in Europe to gain insight into their research practices and interaction with their users. The results show that CFs strive to provide the best possible service to their users. Our results corroborate the ABRF survey’s results, which also found that CFs are deeply invested in implementing best research practices, support transparency, rigor and reproducibility and protect against cognitive bias (Knudtson et al. 2019). The ABRF survey identified the lack of training, mentorship or oversight as the main factors contributing to the lack of compliance with rigorous and reproducible research. Similarly, respondents to our survey cited training, advising and communicating with users as the most important factors for achieving research quality. In both surveys, respondents listed mostly identical tools to improve research quality. On the other hand, the major challenges in promoting best scientific practices differed in the two surveys. While funding and lack of staff was most critical for our respondents, it was poor sample quality and lack of training in the ABRF survey (Knudtson et al. 2019).

Our survey reveals several weaker areas with potential for improvement. Insufficient funding remains the major issue for the majority of CFs and affects not only the ability to purchase and maintain the state-of-the-art equipment, but perhaps even more importantly it also directly affects securing enough qualified staff, training staff and users, and implement a good facility management system. Furthermore, the lack of funding prevents CFs from developing and improving their techniques or from providing a more extended service to the users even if they would like to (Figure S4).

The majority of CFs recognise the need for monitoring the quality through the whole experimental process. Based on the responses we propose that the CFs incorporate at least the following four quality checkpoints to efficiently ensure research quality with the active help of the users: checking the experimental design and 2) the sample quality would avoid running costly experiments unnecessarily and ensure solid data for interpretation. 3) A data analysis check would ensure rigor and transparency and would decrease experimental bias. 4) A final check before publication would allow to make sure the results are presented optimally and comply with best research practices (figure 7). The proposed check points need to be customized to the individual CF needs. For example, blinding and randomization are very important aspects of experimental design in animal core facilities. In addition, only a precise and relevant documentation can guarantee data traceability. All these aspects should be considered to achieve rigor, reproducibility and traceability. Users of self-service facilities would particularly benefit from CF staff expertise as most of CFs offer a lot of information to the users from the very beginning (experimental design) to the end (publication). This is especially relevant for techniques, which are new not only to users but also to their PIs.

**Fig 7.**
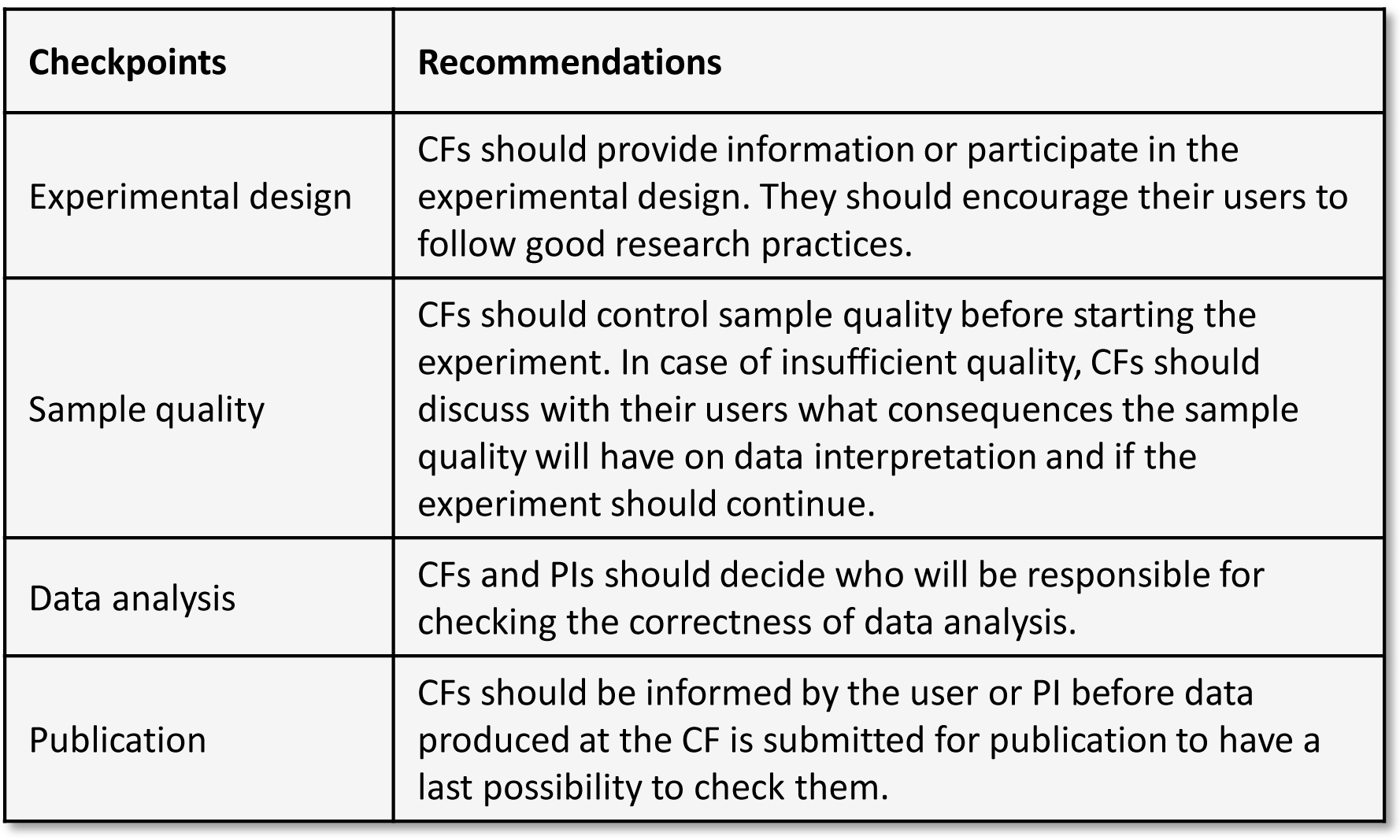
Four important checkpoints for quality.

A management software was rated as one of the best tools to improve research quality. A well-functioning management software also ensures the traceability of produced data. The software solutions used by CF are very heterogeneous with the implementation and cost being the biggest obstacles preventing wider usage. Importantly, some respondents noted that there is currently no management software allowing the full management of the facility, from booking scheduling, experimental design, data acquisition and analysis to billing. Development of such software would likely reduce the time CFs lose by switching between two or more incompatible software packages and increase the traceability of the data. Critically, an ideal management software must be user-friendly, simple and fast, as users are not willing to use an overly complicated and time demanding software and might refuse to cooperate.

Another sensitive point in the CF-user interaction turned out to be also the most fundamental one: communication. Communication between users and CF staff can be challenging as users are not always ready to read or listen to the staff advice, which can directly affect science quality. One third of CFs is not satisfied with the current situation and wishes to improve the communication with their users. This might seem trivial, however, handling questions or requests even from a moderate number of users on an individual basis can easily overload the often-limited CFs’ staff. While the solutions need to be tailored to particular CFs, a good CF management software can lighten the load on the CF staff. Other suggestions were the use of online chats or blogs, with the advantage to directly interact with multiple users at once. Obviously, communication also requires an active participation from the users. Tensions were frequently reported, when users ignore or do not make optimal use of the information that is offered to them by CFs. We cannot fully analyse the reasons for this based on this survey. The CFs should make sure that the provided information including protocols is clear, comprehensive, easy to follow and timely. However, one of the underlying reasons users do not follow the CFs protocols might be ill-defined responsibility.

The involvement of the CF staff in the user’s research raises the question of responsibility for the produced data. CFs have very diverse opinion on the matter, most of them believing that users and PIs are fully responsible for the work. Interestingly, most CFs also think that being part of the publication process would improve the quality of the published report, as they could check the data quality and interpretation. However, this would make them also responsible. Users and PIs bearing the full responsibility for produced data might be relevant in the self-service facilities but cannot be easily justified in the full-service or hybrid CFs. Some CFs do take responsibility for the data, depending on how much involved they were. Along the same lines, the Core for Life, an alliance of Life Science Core Facilities in Europe (www.coreforlife.eu), is aiming, among other, to develop a system to trace and evaluate standard procedure contributions of CF in publications. “This system would clarify responsibilities for the quality of the data generated in the research infrastructures”. It is probably difficult to define a simple universal rule. In any case, the responsibility should be clearly defined before the start of a project, perhaps in a similar way as it is stipulated in the Good Clinical Practice that must be followed in clinical trials (https://www.ich.org/page/efficacy-guidelines). An assignment of responsibility should be an essential part of the user agreement. It should also include an agreement on the authorship and with it associated responsibilities from both sides, the CF and the user.

In summary, the survey highlighted issues that affect the quality of research at CFs and could be remedied by rather simple measures. First, relevant quality checkpoints should be introduced at sample submission, after data analysis and also just before publication. Second, data management should be further improved in most CFs, and the use of a management software would be beneficial. Third, there is a need for improvement of the communication between CF staff and users, which requires tact and effort and is often perceived as a challenge. In addition, the responsibilities of each party should be clearly defined. The presented survey is part of the Q-CoFa (Quality in Core Facilities) project that aims to develop a framework for the best quality practices at the interface between CFs and researchers and provide guidance on communication, information flow and data management to ensure the generation of rigorous and reproducible data. We are working on guidelines which we will publish to further strengthen the role of CFs to increase and promote research data quality.

## Limitations

1000 CFs were invited to participated in the survey and only a quarter completed the survey. It is possible that more quality concerned facilities were more likely to participate in the survey, therefore causing a selection bias. Additionally, the survey targeted only the CF staff and thus it lacks the users’ point of view.

## Material and methods

We developed a 68-question online survey using Limesurvey software and sent it via emails to 1000 Core Facilities in different life science fields throughout Europe. The Core facilities and emails were retrieved using Google search engine or were reached by publicizing our survey in the CTLS newsletter (Core Technologies for Life Sciences). The survey was open from December 2019 to July 2020. All survey participants were anonymous. The survey contained yes/no, multiple-choice and open-field text questions. The survey data was analysed using Microsoft Office 365 Excel. Open-field text questions were evaluated by reading each of them personally and defining categories based on the responses. Keywords were then chosen to allow automatic counting in Excel. The survey questions can be downloaded as Supplementary figure S5. The Excel file with the raw and analysed data can be obtained from the corresponding authors.

## Acknowledgment

We are grateful to Dr. Martin Kos and Dr. Nicolas Sylvius for helpful discussions and critical reading of the manuscript. We thank Dr. Anton Bespalov and Dr. Christoph Emmerich (PAASP) and Dr. Barbara Hendriks (DZHW, Deutsches Zentrum für Hochschul-und Wissenschaftsforschung) for their help designing the survey, valuable discussions and critical reading of the manuscript. We are grateful to Dr. Charles Girardot for helpful discussions about management software. We also thank all the participants of the survey for their time. This work was supported by the Federal Ministry of Education and Research (BMBF, grant 01PW18001, project Q-CoFa) and was performed at the Interdisciplinary Neurobehavioral Core facility at the University of Heidelberg.

## Notes

### Competing Interest Statement

The authors have declared no competing interest.

