## Supplemental Figures 1 to 5 for "Survey of Core Facilities shows the importance of communication and management for optimal research quality"

### Fig S1. Characteristics of the core facilities

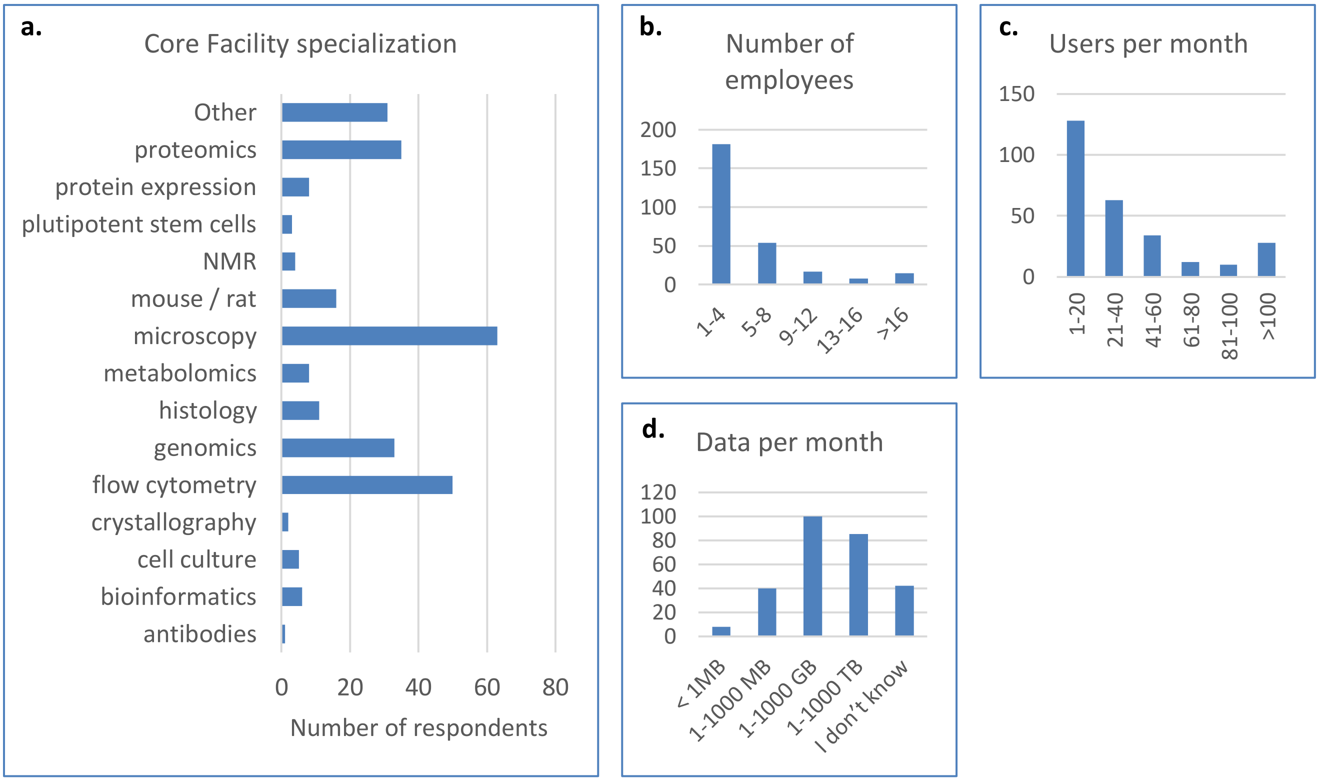


**Figure S1.** Characteristics of the surveyed core facilities.
a) Type of technology provided. NMR, nuclear magnetic resonance.
b) Number of employees,
c) Number of users per month
d) Amount of data produced in a month

### Fig S2. Full-service versus self-service

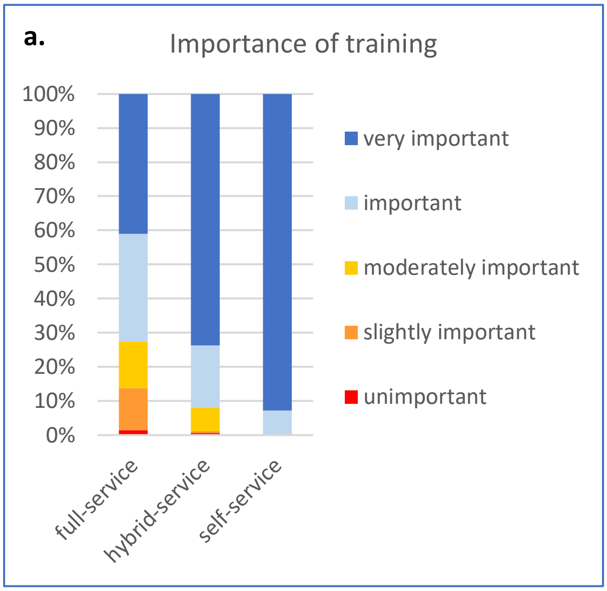


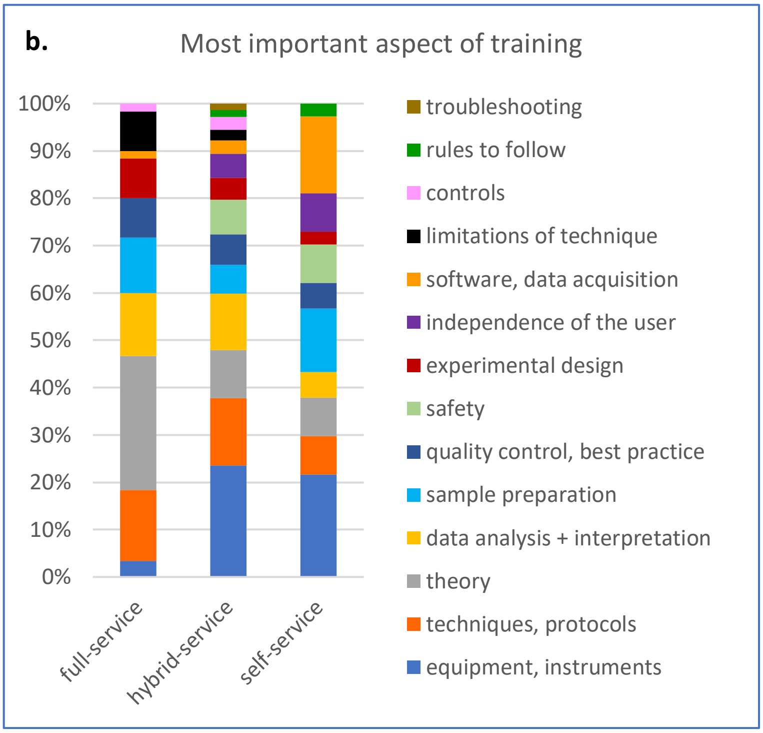


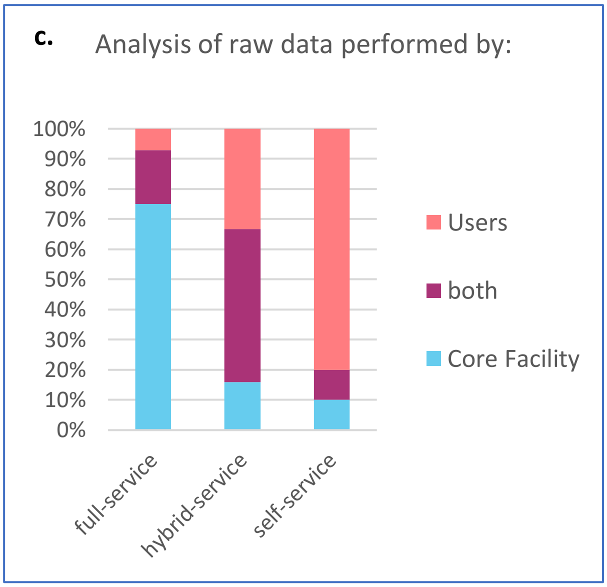


**Figure S2.** Full-service versus self-service facilities.

a) Overall importance of training
b) The most important aspects of training
c) Responsibility for raw data analysis

### Fig S3. Management

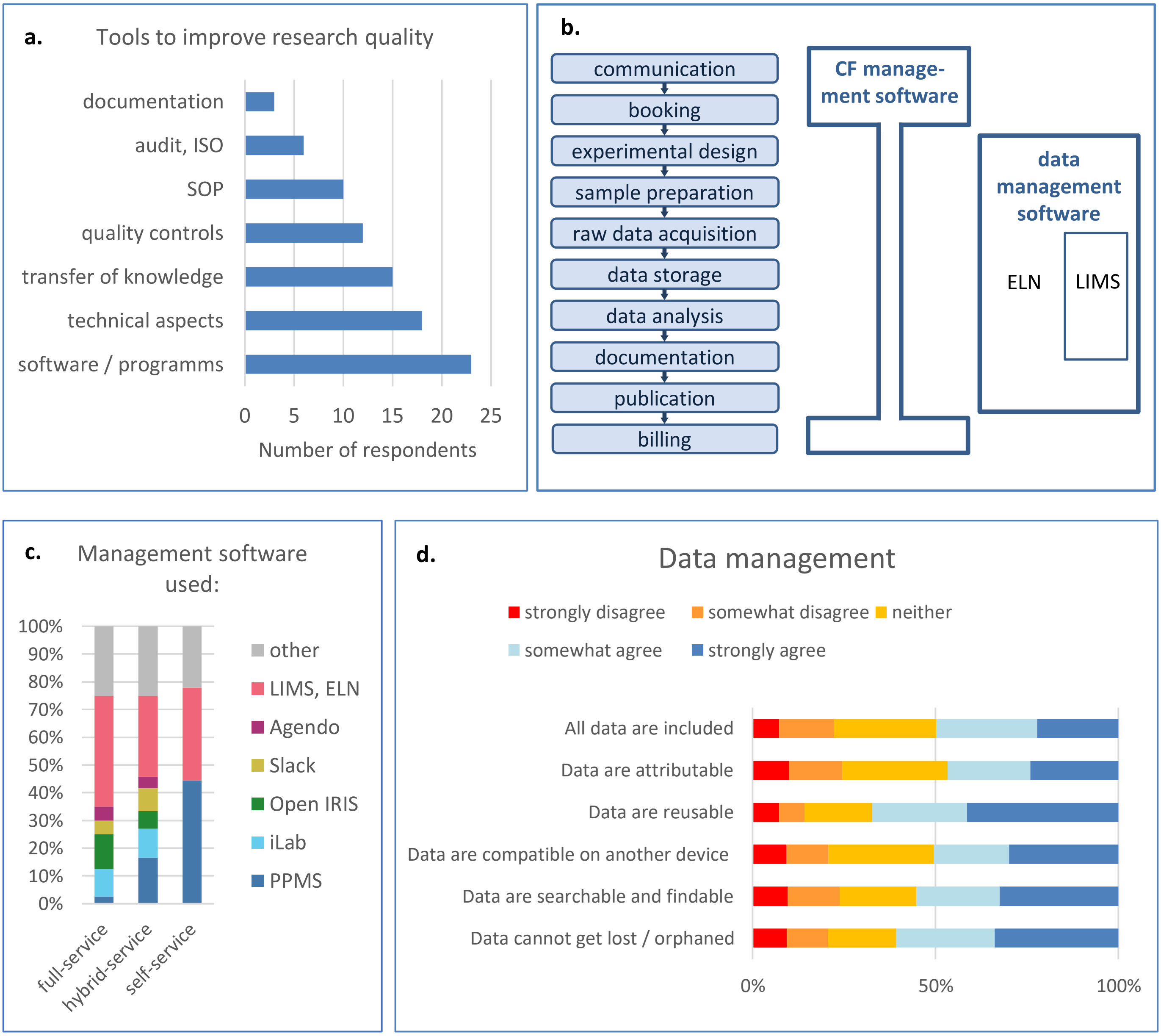


**Figure S3.** Management.
a) Tools to improve research quality. Technical aspects covers instrumentation, robots, automation and improving techniques.
b) Two categories of management software cover different tasks in CFs.
c) Management software used in different CF types
d) Current situation in CFs regarding data management

### Fig S4. Repercussions of insufficient funding on quality

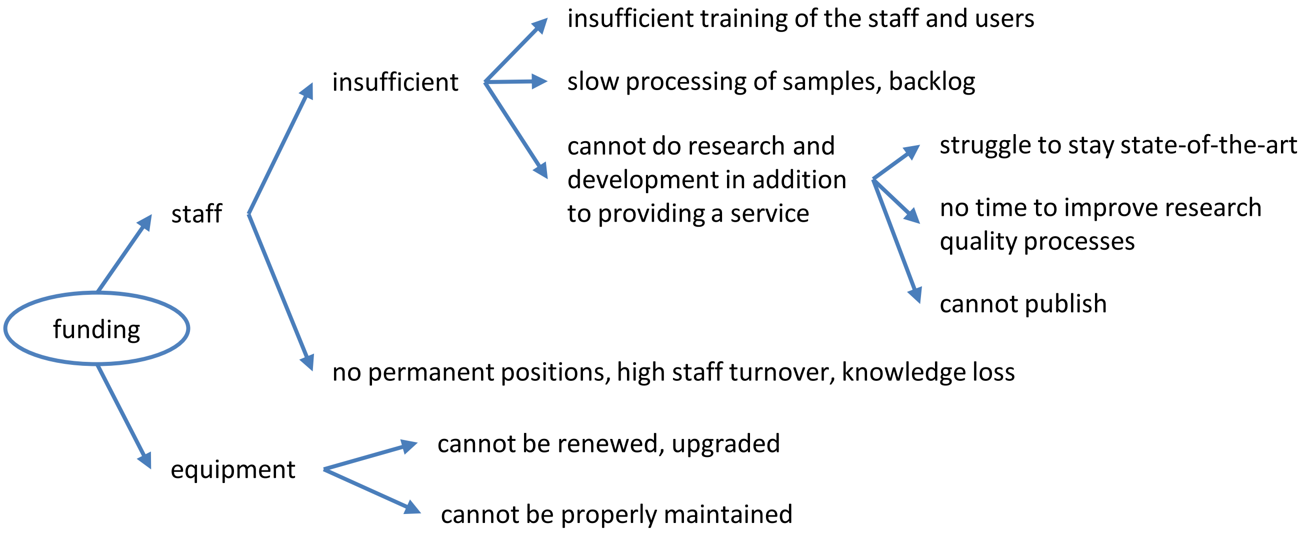


**Figure S4.**
Potential repercussions of insufficient funding on quality

#### Figure S5: Survey questions.

1) **Scope of your core facility**

   a) What is your core facility specialized in?
   antibodies / cell culture / crystallography / flow cytometry / genomics / histology / metabolomics / microscopy / mouse / NMR / pluripotent stem cells / proteomics / protein expression / other (free text)

   b) Who performs the experiments at your core facility?
   The staff of the core facility does (the users don't work at the core facility).
   The users work at the core facility (you provide the infrastructure only).
   Both are possible.

   c) What amount of data is produced per month?
   < 1MB / 1-1000 MB / 1-1000 GB / 1-1000 TB / > 1 PB / I don't know

   d) How many employees do you have? 1-4 / 5-8 / 9-12 / 13-16 / >16

   e) How many users are coming to your core facility per month?
   1-20 / 21-40 / 41-60 / 61-80 / 81-100 / >100 users per month

2) **Communication**

   a) How are you communicating with your users?
   in person, email, phone, website, seminars/workshops, user group meeting, social media, communication/management/scientific software, other (free text)

   b) Would you like to have the communication with your users improved? Yes / no

      i) If "Yes": How? (free text)

   c) Are you using a communication software? Yes / no

   If "Yes":

      i) Which software are you using for communicating with your users? (free text)

      ii) What is your experience? (free text)

   If "No":

      iii) Do you think it would make sense to use a communication platform? Yes / no

      iv) Would you see obstacles? (free text)

3) **Training**

   a) Do you train your users about experimental techniques used in your facility? Yes/no/NA

      i) If "Yes": What is the most important aspect of your training? (free text)

   b) How important do you consider training? unimportant / slightly important / moderately important / important / very important

4) **Experimental design**

   a) Are you involved in experimental design?
   No / Yes, we provide information / Yes, we are actively involved

      i) If "No": Why are you not involved? (free text)

      ii) If "No": Would it be important for you to be involved? Yes/no

      iii) If 1st „Yes": How do you provide it?
      face to face discussion / phone / in writing (file or paper) / email / other (free text)

      iv) If 2nd „Yes": How do you get involved?
      face to face discussion / phone / in writing (file or paper) / email / other (free text)

#### 5) Sample preparation

**a)** Do you provide information to the user on sample preparation? yes / no

i) If "No": Would that be important for you to provide? yes / no

**b)** Do you know what controls the user includes? never / rarely / sometimes / often / always

**c)** Do you know how the user prepares the samples? never / rarely / sometimes / often / always

**d)** If you are aware of poor sample quality do you allow the user to proceed? never / rarely / sometimes / often / always

#### 6) Experimental procedure

**a)** Do you provide written standard experimental protocols? Yes/no/NA

**b)** Do you provide guidance how to increase quality and robustness of research outcome? Yes/no

**i)** If yes: What does it look like? (free text)

**c)** Do you help the users troubleshoot? yes / no

**i)** If "No": Do you think this would be important? yes / no

**d)** Please rate the **current** situation and the **importance** to implement the feature.

1 = strongly agree
2 = somewhat agree
3 = neither agree nor disagree
4 = somewhat disagree
5 = strongly disagree

| Description | **Current situation** | **Importance** |
|---|---|---|
| Samples are randomized. | 1 .... 2 .... 3 .... 4 .... 5 | 1 .... 2 .... 3 .... 4 .... 5 |
| Samples/experiments are replicated. | 1 .... 2 .... 3 .... 4 .... 5 | 1 .... 2 .... 3 .... 4 .... 5 |
| Experiments are performed blinded. | 1 .... 2 .... 3 .... 4 .... 5 | 1 .... 2 .... 3 .... 4 .... 5 |

#### 7) Research data

**a)** Who stores the raw data (long term)? the Core Facility / the user / both / other (free text)

**i)** If "user", "other": Do you think the core facility should also be responsible for storing the raw data? Yes/no (+ free text)

**b)** If CF/both/other: Where are the raw data stored (long term)?
at the core facility physically (PC, external drive…)
internal server/cloud at your organisation/university
external server/cloud through the internet
public repository
other (free text)

**c)** How are the raw data transferred from the core facility to the user?
network (server, cloud, FTP…)
external disk / USB stick
email
other(free text)

**d)** Who performs the primary analysis of raw data? The core facility / the user / both

**i)** If "user/both": Do you provide guidance or train the user how to analyse raw data? Yes/no

**e)** If applicable, who performs the statistical analysis and is that person appropriately trained?
the Core Facility / the user / the core facility and/or user / other (free text)

i) If "user" + "both": Do you provide guidance or train the user how to perform the statistical analysis? Yes/no (+ free text)

f) Is there a quality control mechanism in place to check whether the data was analysed and interpreted correctly? Yes/no

i) If "Yes": Could you please briefly describe? (free text)

ii) If "No": Do you think it would be important to have such quality control mechanism at your core facility? yes / no (+ free text)

g) Do you have a system in place allowing the identification of raw data behind a published figure (where is the raw data stored)? Yes/no

i) If "Yes": Could you please briefly describe? (free text)

h) Are the data backed up? Yes/no

i) Are the data archived? Yes/no

j) Please rate the **current situation** in respect to research data produced in your facility and the **importance to improve** it.

1 = strongly agree
2 = somewhat agree
3 = neither agree nor disagree
4 = somewhat disagree
5 = strongly disagree

| Research data… | **Current situation** | **Importance to improve** |
|---|---|---|
| …cannot get lost / orphaned | 1 …. 2 …. 3 …. 4 …. 5 | 1 …. 2 …. 3 …. 4 …. 5 |
| … are searchable and findable | 1 …. 2 …. 3 …. 4 …. 5 | 1 …. 2 …. 3 …. 4 …. 5 |
| … are compatible / interoperable on another device/software | 1 …. 2 …. 3 …. 4 …. 5 | 1 …. 2 …. 3 …. 4 …. 5 |
| … are reusable | 1 …. 2 …. 3 …. 4 …. 5 | 1 …. 2 …. 3 …. 4 …. 5 |
| … are attributable<br>(Understand who acquired and/or edited the data and when?) | 1 …. 2 …. 3 …. 4 …. 5 | 1 …. 2 …. 3 …. 4 …. 5 |
| … are complete and all data included (any repeat or reanalysis performed on the sample) | 1 …. 2 …. 3 …. 4 …. 5 | 1…. 2 …. 3 …. 4 …. 5 |

#### 8) Documentation

a) Who maintains the lab book/the documentation? the Core Facility / the user / both

b) Do you believe that there is enough documentation about the overall experimental procedure? Yes / no / I don't know

i) If "No": How could that be improved? (free text)

#### 9) Publication

a) Do you provide support for writing the "materials & methods" section for a publication? Yes/no

b) Do the users contact you when writing their publication? never / rarely / sometimes / often / always

c) Do you think it could improve the quality of published research data to involve the core facility in the publication and why? Yes / No (+ free text)

**10) Quality improvement**

a) Do the users actually use the information you provide? never / rarely / sometimes / often / always

    i) What do you do to encourage them to use it? (free text)

b) Do you have a data management plan in place (= a written statement/protocol on the management of research data)? Yes/No

    i) If "Yes": If you are willing to share with us, please contact us.

    ii) If "No": Would you find it useful to implement one and why? Yes / No (+ free text)

c) Do you have enough staff to run your facility? Yes/No

d) Do you have enough funding to maintain and upgrade your equipment? Yes/No

e) Are you currently using any tools to achieve higher research quality in your core facility? Yes/No

    i) If "Yes": What are these tools? (free text)

f) What are the most important factors for achieving research data quality in your core facility? (free text)

g) Where do you see the greatest need for improvements in your Core Facility? (free text)

h) What challenges do you see? (free text)
